# Increases in Plasma Viscosity Disrupt Microvascular Flow Dynamics

**DOI:** 10.1101/2025.11.21.689861

**Authors:** Cristian E. Franco, Albert L. Gonzales

## Abstract

**Background:** Autoregulation of blood flow depends on pressure induced constriction and flow mediated dilation. These processes are well defined in arterioles, but how blood enters and distributes within the capillary network, the primary site of oxygen and nutrient exchange, is less understood. Endothelial cells and pericytes are increasingly recognized as active regulators of capillary tone, yet the mechanisms governing capillary level autoregulation remain unclear. Plasma viscosity is a major determinant of shear stress and may influence this regulation, but its role is not fully defined.

**Methods:** Using an *ex vivo* pressurized retinal preparation that preserves the arteriole to capillary continuum, we examined how changes in intraluminal viscosity and shear stress affect endothelial and mural cell activity during pressure driven flow. To study chronic viscosity elevation, we assessed microvascular structure, remodeling, endothelial responsiveness, and capillary perfusion in a high fat diet model of altered blood rheology.

**Results:** Increasing viscosity enhanced endothelial Ca^2+^ activity in transitional and capillary segments and suppressed pressure induced Ca^2+^ elevations in smooth muscle cells and pericytes through nitric oxide dependent inhibition. Chronic elevation of viscosity produced segment specific effects. Arterioles showed structural remodeling and higher tone, whereas transitional and capillary vessels did not remodel and instead exhibited impaired endothelial shear sensing and reduced capillary recruitment. High fat diet feeding lowered baseline capillary perfusion and eliminated viscosity dependent modulation, indicating loss of shear-based control.

**Conclusions:** Autoregulation differs fundamentally between arterioles and capillaries, and viscosity dependent endothelial signaling is a key determinant of capillary entry and flow distribution. Plasma viscosity is an important regulator of microvascular perfusion and a potential biomarker and therapeutic target in microvascular disease.

## Introduction

Vascular autoregulation enables blood vessels to maintain constant blood flow despite changes in blood pressure, ensuring that cells receive the O_2_ and nutrients required for proper function. In resistance arteries and arterioles, endothelial and smooth muscle cells respond differently to mechanical forces: endothelial cells promote dilation in response to increased flow, while smooth muscle cells induce constriction in response to increased intraluminal pressure. Traditionally, it was believed that the capillary network, comprising capillary endothelial cells and pericytes, lacked intrinsic autoregulatory mechanisms, with blood flow regulation primarily attributed to upstream arterioles and precapillary sphincters (1, 2). However, recent studies performed in the brain and retina have challenge this perspective, demonstrating that capillary endothelial cells and pericytes form an active sensory network that detects neural activity and regulates local blood flow (3–5). In addition, Klug et al. (6) and Ferris et. al. (7) demonstrated that distinct pericyte subtypes, ensheathing pericytes in the transition zone and thin-stranded pericytes deeper in the capillary bed, both exhibited Ca^2+^ elevation and sustained contraction in response to increased intraluminal pressure, a response necessary to maintain cerebral capillary blood flow. This response allows them to act as pressure-sensing autoregulators of blood flow within the capillary network, helping to prevent hypoxia and endothelial damage (6–8). To deepen our understanding of microvascular autoregulation, it is critical to investigate how other mechanical forces, such as fluid flow and shear stress, contribute to blood flow regulation within capillary networks.

Shear stress, the frictional force generated by blood flow along the endothelial surface, is a key contributor to vascular autoregulation. In larger vessels such as arteries and arterioles, shear stress influences endothelial signaling, vascular tone, and structural remodeling (9). These effects are tightly coupled to the non-Newtonian, shear-thinning properties of blood, which allow viscosity to decrease with increasing flow velocity (10). While the mechanotransduction of shear stress in larger vessels has been extensively studied, far less is known about how shear forces are sensed and transduced in the capillary network. Capillaries represent the primary site of O_2_ and nutrient exchange and exhibit unique flow dynamics, yet the role of shear stress in capillary endothelial signaling and pericyte-mediated microvascular regulation remains poorly defined. In this study, we show that intraluminal viscosity strongly influences how blood enters and distributes within the capillary network by differentially regulating endothelial and mural cell Ca^2+^ signaling. Increased viscosity enhances endothelial Ca^2+^ activity while simultaneously suppressing pressure-induced Ca^2+^ elevations in arteriolar smooth muscle cells and pericytes through nitric oxide–dependent mechanisms.

Notably, chronic elevation of plasma viscosity induces structural remodeling and increased tone in arterioles, whereas capillary and transition segments do not remodel and instead exhibit impaired shear sensing and reduced capillary perfusion. Together, these findings demonstrate that autoregulatory mechanisms at the capillary level differ fundamentally from those in arterioles and identify viscosity-driven endothelial signaling as a key determinant of capillary entry, microvascular perfusion, and tissue health.

## Methods

### Animals

Animals were housed and treated according to protocols set by the Institute Animal Care and Use Committee (IACUC approval no. 20-07-1039-1) at the University of Nevada, Reno. Adult (2–5-month-old) male and female mice were group-housed with environmental enrichment and free access to food and water. C57BL/6J mice were used as controls. *Myh11*-GCaMP6f-Ai95D, *Cspg4*-GCaMP6f and *Cdh5*-GCaMP6f-Ai95D transgenic mice were used for the cell-specific expression of Ca^2+^ indicator GCaMP6f in vascular mural cells and endothelial cells, respectively. *Myh11*-GCaMP6f and *Cspg4*-GCaMP6f mice (6) are generated by crossing mural cell-specific myosin heavy chain 11 (*myh11)*-Cre (037658; Jackson Laboratory, Bar Harbor, ME) or chondroitin sulfate proteoglycan 4 (cspg4)-Cre mice (008533; Jackson Laboratory, Bar Harbor, ME) with floxed high signal-to-noise Ca^2+^ indicator GCaMP6f mice (028865; Jackson Laboratory, Bar Harbor, ME). *Cdh5*-GCaMP6f-Ai95D mice (11) are generated by crossing cadherin5 (*cdh5)*-Cre mice (006137; Jackson Laboratory, Bar Harbor, ME) with floxed GCaMP6f mice (028865; Jackson Laboratory). To induce GCaMP6f expression, mice were fed tamoxifen-containing chow (0.25g/kg, inotiv) for two weeks immediately after weaning, then returned to standard chow for an additional two weeks to allow for transgene recombination and stabilization of expression. To model chronically elevated plasma viscosity, mice were fed a non-obesogenic high-fat diet (1% cholesterol, 15% calories from fat; TD.09237, inotiv) for three months beginning at weaning.

### Solutions and Reagents

Retina dissection buffer consists of 119 mM NaCl, 3 mM KCl, 0 mM CaCl_2_, 3 mM MgCl_2_, 5 mM glucose, 26.2 mM NaHCO_3_ and 1 mM NaH_2_PO_4_, bubbled with 95% O_2_/5% CO_2_ (pH 7.4). Retinal physiological saline solution (PSS; pH 7.4) contains 124 mM NaCl, 26 mM NaHCO_3_, 1 mM NaH_2_PO_4_, 2.5 mM KCl, 1.8 mM CaCl_2_, 2 mM MgCl_2,_ and 10 mM glucose, bubbled with 95% O_2_/5% CO_2_. Mg^2+^-supplemented PSS contains 124 mM NaCl, 26 mM NaHCO3, 1 mM NaH_2_PO4, 2.5 mM KCl, 3.8 mM MgCl2, and 10 mM glucose. All chemicals and drugs were purchased from Sigma-Aldrich (St. Louis, MO, USA) unless specified otherwise.

### *Ex Vivo* Retinal Pressurization Preparation

Animals were deeply anesthetized using isoflurane 5% and euthanized by exsanguination and decapitation. As previously described in Klug et al.(6), for *ex vivo* pressurized retina preparations, the entire orbital structure—including the eye, optic nerve, ophthalmic artery, and surrounding musculature—was carefully excised and transferred into ice-cold, Ca^2+^-free, Mg^2+^-supplemented rPSS bubbled with 95% O_2_ / 5% CO_2_. Extraneous outer tissues, such as muscles and nerves, were trimmed away using fine scissors, leaving the ophthalmic artery, optic nerve, and associated fine musculature intact. Branches of the ophthalmic artery not supplying the central retinal artery were ligated with fine suture. For pressurized preparations, the dissected retina was transferred to a custom myography chamber, and the ophthalmic artery was cannulated using a glass micropipette filled with rPSS and mounted on a micromanipulator. To allow the retina to lie flat, strategic incisions were made along the retinal edge, enabling it to spread on a custom silicon platform with the cannulated artery and surrounding structures positioned within the imaging plane. The retina was continuously superfused with 37 °C rPSS at a rate of 10 mL/min. Intraluminal pressure was established by connecting the cannula to a gravity-fed column and a perfusion pressure monitor (PM-4, Living Systems), which was adjusted to ∼60 mmHg to reflect physiological pressure at the ophthalmic artery. A two-way valve was opened to initiate perfusion. Successful cannulation and pressurization were confirmed by rapid clearance of blood cells from the retinal vasculature.

### Confocal Imaging

The imaging of Ca^2+^ dynamics, pressure-induced tone, and flow of fluorescent microspheres was performed on a stand-alone Crest Optics X-Light V3 spinning disk confocal and widefield microscope (Rome, Italy) with 7-wavelength laser launch (LDI-7, 89 North), dual ORCA-Fusion Gen-III sCMOS Cameras (Hamamatsu). Retinal vasculature was stained intraluminally by FITC-conjugated isolectin B4 and imaged using a spinning disc upright Olympus confocal microscope using 10x and 60x magnification. The cannula was filled with rPSS and kept at 40mmHg until pressure changes were required. Additionally, the intraluminal solution viscosity was increased by the addition of glycerol. To visualize cell-specific GCaMP6f Ca^2+^ dynamics. The peak excitation (λ_ex_) and emissions (λ_em_) wavelengths of GCaMP6f are 496 nm and 513 nm, respectively were acquired at a resolution of 512 × 512 pixels at 30 frames/s at 37°C. Intraluminal pressure was increased, resulting in increased fluorescent intensity, indicating an increase in intracellular Ca^2+^. Intraluminal viscosity was increased using glycerol to increase shear stress within the vessel. To visualize fluid flow, fluorescent polystyrene microspheres (FSDG005, Bangs Labs) with a 2.07 µm diameter were used. The beads were diluted to 1:500 in 1x PBS with 1% bovine serum albumin to prevent clumping. The solution was then sonicated for 5 min with 5 μL loaded into the cannula, further diluting the solution, at 40 mmHg, and visualized at 10 ms exposure for 100 frames/second. Velocity was measured by measuring the distance covered in μm at each 10-ms exposure frame to determine the average speed per microsphere in μm/ms.

### Image and Statistical Analysis

Images were processed and analyzed using ImageJ and SparkAn software (created by Adrian Bonev and provided by Mark T. Nelson, University of Vermont). For Ca^2+^-imaging experiments, baseline fluorescence was determined by the first 100 frames before the pressure step, divided by the background fluorescence (F_0_), and normalized to F-F_0_. The change in F/F_0_ was used to show a fluorescent increase in *myh11*-CGaMP6f and/or *cspg4*-GCaMP6f mice. The area under the curve was taken to identify peaks higher than 10 times the standard deviation for the baseline to determine the frequency of Ca^2+^ events for *cdh5*-GCaMP6f mice. For microsphere experiments, linescans were run at transition segments’ capillary junctions to track the frequency of fluorescent microspheres. The number of spheres for each experiment was kept consistent by injection of 5 μL of microsphere stock solution. The angle of deviation measurement was measured from the parent branch towards the selected daughter branch.

Data in figures and text are presented as means ± standard error of the mean (SEM). Values of “n” refer to the number of experiments, and “N” refers to the number of animals. Statistical analyses were performed using Prism 10 (GraphPad). Statistical significance was determined using unpaired student’s t-tests, one-way analysis of variance (ANOVA), two-way repeated-measures ANOVA. P-values ≤ 0.05 were considered statistically significant for all experiments.

## Results

### Fluid Viscosity Delays the Filling of Capillary Vessels

Fluid viscosity plays a major role in flow resistance, with higher viscosity increasing resistance and lower viscosity allowing smoother and more efficient flow through a straight tube. To investigate the impact of intraluminal viscosity on fluid behavior within the native capillary microcirculation, a network of interconnected tubes, we used a pressurized retinal vasculature preparation (6, 8, 12) in combination with a high-speed, high-resolution spinning disk confocal microscope. In this preparation, the retina and its attached vasculature were isolated, and the ophthalmic artery was cannulated using a glass pipette. The retina was then positioned on a custom silicone platform, flattened, and secured with short tungsten wire pins (Fig. 1A and B). To alter intraluminal viscosity, we supplemented PSS with 40% glycerol, increasing the cannula physiological saline solution (PSS) viscosity from 1.0 cP to 3.0 cP, respectively.

**Figure 1.**
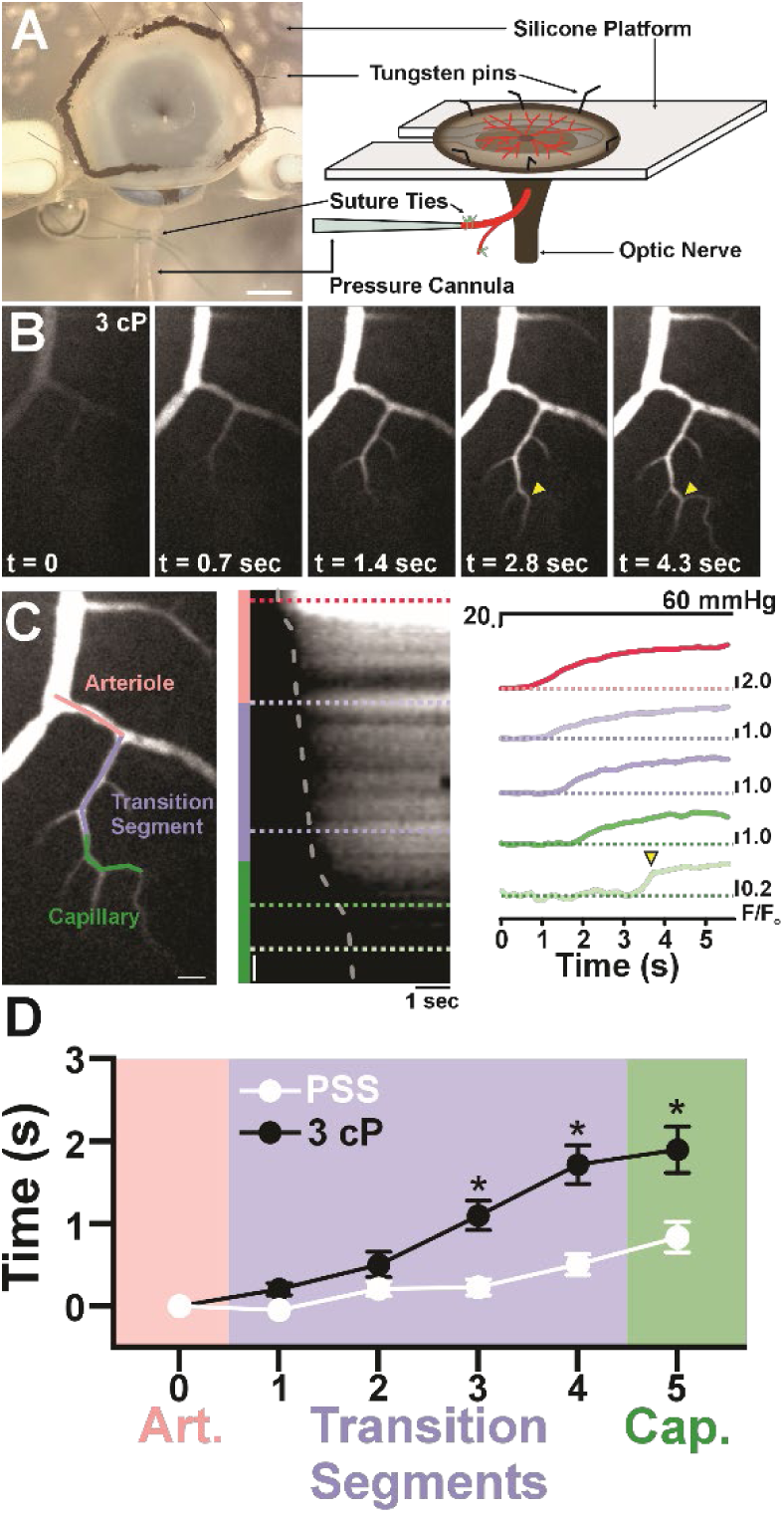
Fluid Viscosity Delays the Filling of Capillary Vessels. **A, Left**: Representative image of an isolated retina mounted on a custom silicone platform and secured with tungsten wire pins. Scale bar = 1 mm. **Right**: Schematic illustrating retinal flattening, positioning, and cannulation of the ophthalmic artery to control inlet pressure. **B**: Representative time-series images showing delayed fluorescent filling of the microvascular network at higher viscosity (3 cP) under identical pressure conditions. **C:** Representative image identifying arterioles, transition segments (1st–4th order), and capillaries (5th order).. Line-scan kymographs and corresponding fluorescence traces demonstrate increasing filling delay in downstream segments during a pressure step. Scale bars = 20 µm. **D:** Summary data showing that increased viscosity (3 cP, black) significantly prolongs filling time compared with PSS/1 cP (white) across arterioles and 1st–5th order branches. Data are presented as mean ± SEM; *p ≤ 0.05 vs PSS; two-way repeated-measures ANOVA with Tukey’s multiple-comparison test; n = 17–19 vessel segments; and N = 3 mice.

Fluorescein was added to visualize fluid distribution (Fig. 1B) from radiating arterioles, transition regions defined as the first four branches with ensheathing pericytes, and distal capillaries characterized by fifth-order and higher branches with thin-stranded pericytes (13). By generating kymographs (Fig. 1C), we measured the fluorescence intensity along a vascular path over time, allowing us to observe filling rates in different branches of the capillary network. To quantify the time to half-maximal fluorescence, we calculated the time delay in fluorescence increase between the radiating arteriole and the first five capillary branches (Fig. 1C). When intraluminal pressure in the ophthalmic artery was increased from 40 to 60 mmHg using 1 cP PSS, filling of the 2nd to 5th order capillaries followed a linear trajectory, occurring on average between 0.21 ± 0.09 seconds and 0.83 ± 0.19 seconds, respectively. When the viscosity of the PSS within the pressure cannula was increased to 3 cP, filling of the 3rd to 5th order capillaries followed a non-linear trajectory, with average times ranging from 0.50 ± 0.15 seconds to 1.9 ± 0.28 seconds. Notably, at several capillary bifurcations, flow in one branch was delayed by seconds (Fig. 1B and C, yellow arrowhead), suggestive of a dynamic opening mechanism permitting downstream flow. These findings demonstrate that increased fluid viscosity alters the timing of capillary filling, particularly in deeper branches, prompting the question of which cellular autoregulatory mechanisms govern flow distribution within the microvascular network.

### Intraluminal Viscosity Increases Endothelial Cell Intracellular Ca^2+^

Fluid viscosity is a key factor in determining shear stress, a force sensed by endothelial cells that line all blood vessels; however, few studies have directly examined its impact on endothelial cell responses. Instead, most research relies on indirect methods, such as increasing intraluminal pressure to increase flow and infer these effects (14, 15). Therefore, to assess the impact of viscosity on endothelial cells’ Ca^2+^ dynamics, we used *cdh5*-GCaMP6f transgenic mice, in which the GFP-based Ca^2+^ indicator, GCaMP6f, is specifically expressed in endothelial cells under the control of the cadherin-5 (*cdh5*) promoter. Due to erythrocyte deformation and aggregation (16), whole blood behaves as a shear-thinning non-Newtonian fluid with a viscosity ranging from 3.5 to 5.5 cP (5). In contrast, plasma is a Newtonian fluid with a viscosity of 1.3 to 1.7 cP (17). To increase viscosity, we supplemented PSS with 30% or 40% glycerol, raising the cannula fluid viscosity from 1.0 cP to 1.6 cP and 3.0 cP, respectively. Using 1.0 cP PSS in the cannula, we elevated ophthalmic artery pressure from 20 to 60 mmHg. The increase in intraluminal flow triggered multiple transient spikes in endothelial Ca^2+^ activity throughout the arterioles, transition segments, and capillaries (Fig. 2A). When intraluminal viscosity was increased to 1.6 cP and 3.0 cP, the pressure-induced increase in flow significantly enhanced the Ca^2+^ response in transitional and capillary endothelial cells. The higher viscosity induced a robust and sustained Ca^2+^ signal (Fig. 2A) that persisted until the intraluminal pressure was reduced. These findings demonstrate that increased intraluminal viscosity converts and amplifies endothelial Ca^2+^ responses to pressure-induced flow, particularly in transitional and capillary endothelial cells. The enhanced and prolonged Ca^2+^ signaling at higher viscosities highlights the sensitivity of endothelial cells to mechanical cues, with a more pronounced attenuation of transient signals observed at 3.0 cP, suggesting viscosity-dependent modulation of endothelial mechanotransduction.

**Figure 2.**
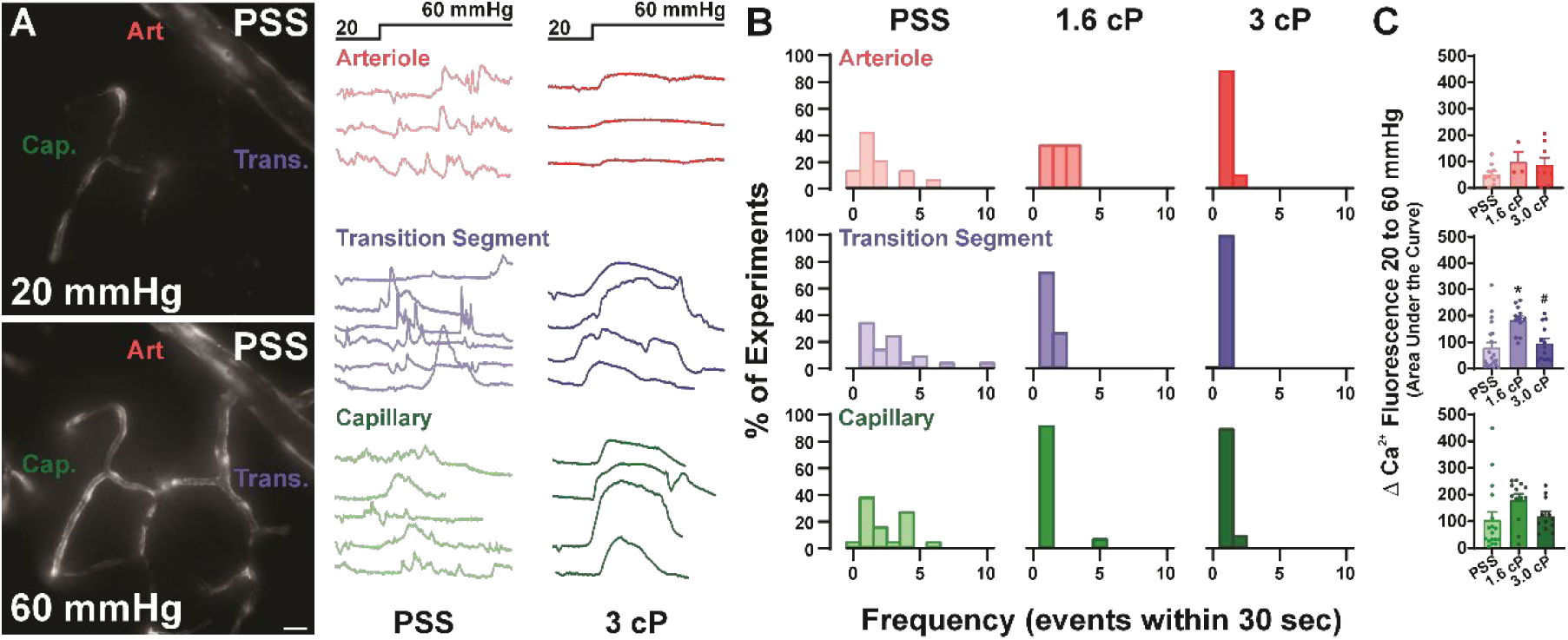
Intraluminal Viscosity Converts Capillary Endothelial Ca^2+^ Signals. **A:** Representative image and traces showing *cdh5*-GCamp8f signaling from pressure increase from 20-60 mmHg. Scale bar = 20 µm**. B:** Histogram summary of fluorescent increase from pressure increases due to increased viscosity, showing increased fluorescence in transition and capillary segments with increased viscosity. 1cP shows increased transient Ca^2+^ signals, 1.6cP decreased transient Ca^2+^ signal, and 3cP shows a sustained Ca^2+^. **C:** Summary data showing pressure-induced changes in Ca^2+^ signaling in arterioles (top), transition segment (middle), and capillary segments (bottom). Data presented with means ± SEM; *p ≤ 0.05 vs PSS and #p ≤ 0.05 vs 1.6 cP; one-way ANOVA with Tukey’s multiple comparison test; n = 7-23; and N = 3-6 mice.

### Intraluminal Viscosity Decreases Mural Cell Ca^2+^

Shear stress plays a critical role in modulating myogenic tone (18–20) by activating endothelial cell signaling pathways that counteract pressure-induced contractions and relax the surrounding vascular smooth muscle cells. Triggered by an increase in cytosolic Ca^2+^ and the activation of endothelial nitric oxide synthase (eNOS), nitric oxide (NO), a key vasorelaxant factor, diffuses to vascular smooth muscle cells, where it inhibits multiple components of pressure-induced vasoconstriction (15). To study the impact of viscosity on the cytosolic Ca^2+^ of mural cells (SMC and pericytes), we used retinas isolated from *myh11*-GCaMP6f or *cspg4*-GCaMP6f transgenic mice, in which the GFP-based Ca^2+^ indicator, GCaMP6f, is specifically expressed in mural cells under the control of either the myosin regulatory light chain 11 (*myh11*) or chondroitin sulfate proteoglycan 4 (*cspg4)* promoters. Using 1 cP PSS in the cannula, we increased intraluminal pressure from 20 mmHg to 60 mmHg, resulting in a rise in mural cell Ca^2+^ levels, consistent with previous reports on the mechanism of pressure-induced tone (6). When viscosity was increased to 1.6 cP and 3.0 cP, pressure-induced elevations in cytosolic Ca^2+^ were reduced in vascular smooth muscle cells and capillary pericytes (Fig. 3). Additionally, the loss of pressure-induced Ca^2+^ responses measured under 3 cP conditions was restored to near 1 cP levels following nitric oxide synthase (NOS) inhibition with L-NAME (Nω-Nitro-L-arginine methyl ester, 100 µM), suggesting that the reduced Ca^2+^ signaling in mural cells was due to NO-mediated inhibition of their contractile response. These findings indicate that increased luminal viscosity modulates the pressure-induced Ca^2+^ responses, likely through NO-mediated inhibition of mural cell constriction in vascular smooth muscle cells and ensheathing pericytes. In contrast, thin-stranded pericytes maintain their Ca^2+^ responses irrespective of L-NAME treatment, highlighting functional heterogeneity among mural cell subtypes in their response to mechanical and NO signals.

**Figure 3.**
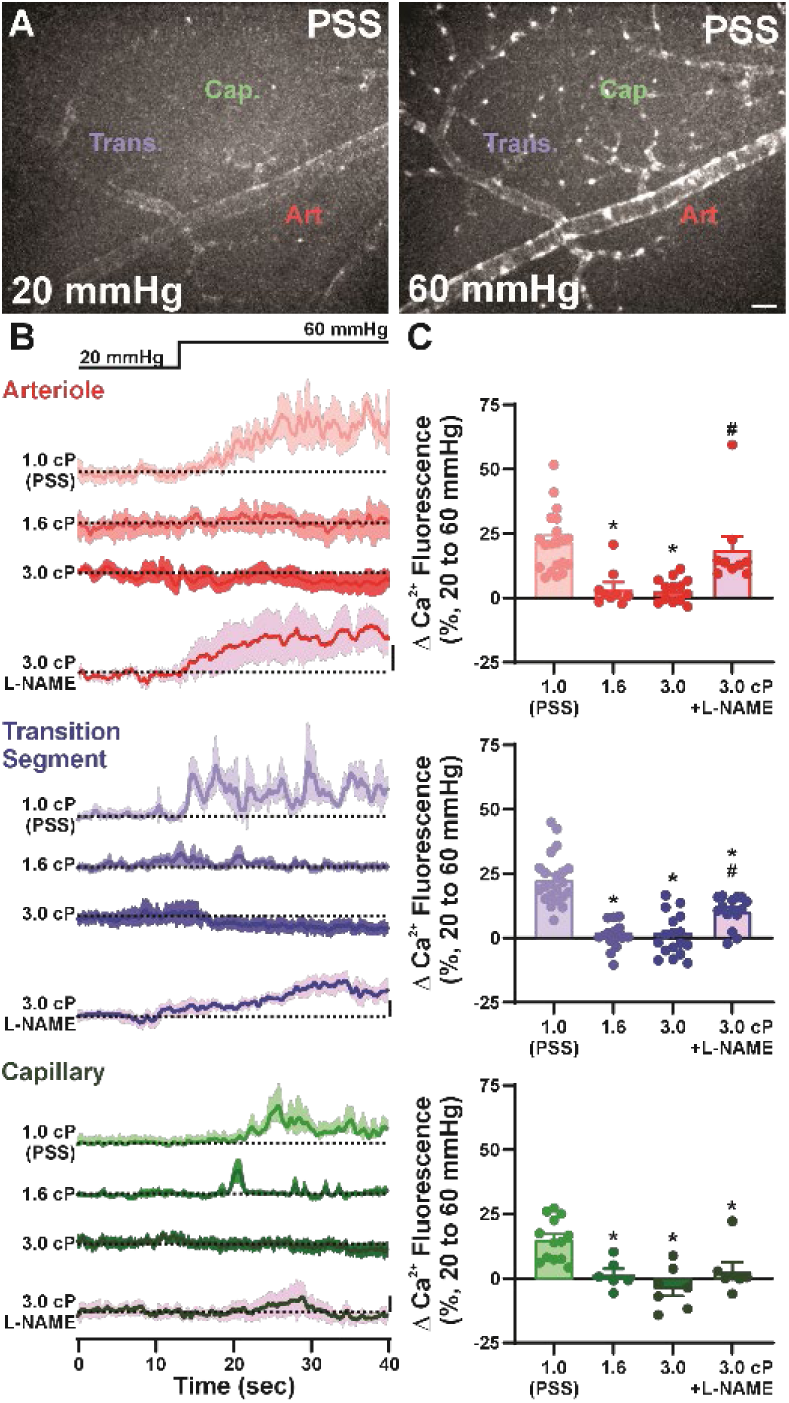
Increased Intraluminal Viscosity Lowers Mural Cell Ca^2+^ Signaling. **A:** Representative fluorescence images of mural cell Ca^2+^ from arterioles (Art.), transition segments (Trans.), and capillary (Cap.) regions at 20 (left) and 60 (right) mmHg intraluminal pressure. Scale bar = 20 µm**. B:** Representative Ca^2+^ fluorescence traces (ΔF/F) from arterioles (top, red), transition segments (middle, blue), and capillaries (bottom, green) recorded during pressure elevation from 20 to 60 mmHg. **C:** Summary data of the pressure(flow)-induced change in mural cell Ca^2+^ at various intraluminal viscosities (1.0, 1.6, and 3.0 cP) and in the presence of nitric oxide synthase (NOS) inhibitor, L-NAME (100 µM). Data presented with means ± SEM; *p ≤ 0.05 vs PSS and #p ≤ 0.05 vs 3.0 cP; one-way ANOVA with Tukey’s multiple comparison test; n=7-20; and N = 4-12 mice.

### High-fat Diet Increases Plasma Viscosity and Impairs Pressure-induced Tone Generation

Transient increases in fluid viscosity influence vascular function by modulating blood flow and pressure-induced tone generation. High-fat diets have previously been shown to alter blood composition and increase plasma viscosity (21). To investigate the chronic effects of elevated fluid viscosity on vascular function, we assessed pressure-induced vasoconstriction by vascular smooth muscle cells and capillary pericytes in mice fed a high-fat diet (HFD) consisting of 1% cholesterol, 15% calories from fat for three months, which did not induce obesity. Plasma samples showed increased viscosity (∼1.6cP) in the HFD mice compared to control mice (∼1.3cP), likely due to the increased circulating triglycerides and cholesterol (21). Previous studies have suggested that high-fat diets (HFD) can impair (22) or enhance (23) pressure-induced tone generation in arteries, potentially due to increased endothelial nitric oxide production (22) or endothelial dysfunction (23). To explore this, we compared pressure-induced tone generation in radiating retinal arterioles, transitional segments (ensheathing pericytes), and capillary vessels (thin-stranded pericytes) from HFD-fed mice and wild-type (WT) mice fed normal chow. Using 1 cP PSS in our pressure cannula, HFD mice exhibited an increased pressure-induced tone generation compared to controls, suggesting an increase in vascular remodeling in the HFD mice. In contrast, no differences were observed in transitional segments or capillaries between groups. When viscosities were increased to 1.6 cP and 3.0 cP, arterioles from HFD-fed mice showed a downward shift in pressure-induced tone generation relative to controls. Notably, while arterioles exhibited the expected viscosity-dependent shift in tone, transitional segments and capillaries from HFD-fed mice showed no changes. These findings suggest that chronic HFD feeding and elevated plasma viscosity promote arteriole remodeling and enhance tone generation, whereas pericyte-covered transitional segments and capillaries do not remodel but instead exhibit a selective impairment in endothelial shear-stress sensing. These findings suggest that chronic high viscosity triggers divergent adaptive strategies in arterioles and capillaries, with arterioles compensating by increasing smooth muscle tone and capillaries reducing their sensitivity to shear stress, thereby shifting overall flow regulation toward greater reliance on endothelial-mediated dilation. These contrasting modes of autoregulation raise important questions about how such mechanisms influence the distribution of blood flow from arterioles into the capillary network.

### Intraluminal Viscosity Decreases Fluorescent Microsphere Distribution

Differences in autoregulation between HFD-induced arteriole and capillary remodeling highlight the need to understand how vascular smooth muscle cells and capillary pericytes use distinct mechanisms to regulate blood flow entering the capillary networks from the feeding arterioles. The superficial retinal vasculature consists of radiating arterioles branching outward from the optic nerve head (Fig. 5A). In proximal regions, capillary networks arise through lateral or orthogonal branching (Fig. 5A), often at ∼90° angles, allowing for uniform perfusion across the retina. Further down the arteriole, branching becomes dichotomous (Fig. 5A), where vessels split into progressively smaller daughter branches, promoting smooth and efficient flow. Previous work has suggested that ensheathing pericytes (1) or capillary sphincters (2) at orthogonal branch points within the post-arteriole transition segments act as “gate-keepers,” regulating blood distribution into the capillary network. Based on this, we tested how transient and chronic increases in viscosity affect fluid entry into capillaries. To address this, we used an *ex vivo* pressurized retinal preparation in which fluorescent beads were introduced through the pressure cannula and tracked as surrogates for intraluminal flow. Kymographs generated from line paths along arterioles and capillaries were used to track fluorescent bead movement, enabling quantification of the fraction entering the capillary network versus continuing through the arteriole (Fig. 5B, right). In control retinas, the fraction of beads entering capillaries declined progressively with increasing viscosity (Fig. 5C). At baseline viscosity (1.0 cP), an average of 39.1% ± 3.9% of beads entered the capillary network. This fraction was significantly reduced at 1.2 cP (23.0% ± 4.3%) and further at 3.0 cP (12.3% ± 4.1%). A smaller but significant reduction was also observed between 1.2 and 3.0 cP. Variability also narrowed at higher viscosities, with bead entry more tightly clustered at lower values compared to the broader distribution at baseline. These results indicate that elevated viscosity reduces both the magnitude and heterogeneity of plasma entry into the capillary network. In contrast, mice fed an HFD exhibited markedly lower bead entry across all viscosities compared to controls, with no significant changes as viscosity increased (Fig. D). Unlike controls, where viscosity-dependent reductions were evident, HFD vessels showed consistently low entry with limited variability, suggesting that capillary perfusion and flow regulation are already impaired. Together, these results demonstrate that viscosity impairs microcirculatory flow by reducing velocity and limiting plasma (bead surrogate) entry into capillaries. HFD further exacerbates this effect by suppressing baseline capillary perfusion, effectively overriding viscosity-dependent partitioning and revealing a compounded mechanism of impaired perfusion under pathological conditions.

## Discussion

In the present study, we used an *ex vivo* pressurized retina preparation to investigate how intraluminal viscosity influences blood flow dynamics and shear stress regulation within the retinal microcirculation. Increasing viscosity delayed capillary filling relative to neighboring arterioles, an effect that was most pronounced at lower intraluminal pressures and diminished at higher pressures, consistent with the physiological compensation of elevated blood pressure under hyperviscous conditions. Elevated viscosity also amplified endothelial Ca^2+^ responses, particularly in transitional and capillary endothelial cells, while reducing Ca^2+^ signaling in vascular smooth muscle cells and capillary pericytes via NO-mediated inhibition. Chronic high-fat diet feeding increased plasma viscosity and promoted arteriole remodeling with enhanced tone generation, whereas transitional segments and capillaries failed to remodel and instead exhibited impaired endothelial shear-stress sensing. Finally, fluorescent microsphere tracking showed that elevated viscosity disrupted the distribution of flow into capillary branches, indicating a loss of controlled flow regulation. Together, these findings demonstrate that both acute viscosity changes and chronic diet-induced hyperviscosity differentially affect endothelial and mural cell function, highlighting viscosity-dependent and cell–type–specific mechanisms that govern blood flow distribution into the capillary network.

### Role of RBCs in Viscosity and Implications for Microcirculatory Flow

Red blood cells (RBCs), though absent in the current experimental model, are essential in determining the rheological properties of blood. Whole blood behaves as a shear-thinning non-Newtonian fluid, meaning its viscosity decreases as shear rate increases. This property allows blood to flow more efficiently under physiological conditions, especially in narrow vessels where shear rates are high (24, 25). In the absence of RBCs, this shear-thinning behavior is lost, which may exaggerate the negative effects of elevated plasma viscosity observed in our experiments. When plasma viscosity rises, microcirculatory flow is reduced, as shown by the delayed capillary filling and decreased intraluminal velocity in our model. *In vivo*, this reduction in flow would likely be even more pronounced, since the presence of RBCs could create a feedback loop: slower blood flow promotes greater erythrocyte aggregation, which in turn further increases viscosity and worsens circulatory impairment. This cycle is particularly concerning in disease states, where blood composition and endothelial responses are already altered. In healthy individuals, compensatory mechanisms such as shear-stress–mediated vasodilation often prevent increased viscosity from raising blood pressure. However, in pathological conditions, these mechanisms are impaired, and clinical studies consistently show that elevated blood viscosity correlates with higher blood pressure and vascular dysfunction (19, 26). Thus, while our findings reveal the impact of plasma viscosity alone, the contribution of RBCs *in vivo* would be expected to further magnify the hemodynamic consequences of increased viscosity within the microcirculation.

### Endothelial and Pericyte Signaling in Pressure-Flow Autoregulation

Increased intraluminal viscosity attenuated pressure-induced tone, as shown by reduced constriction (Fig. 4) and diminished Ca^2+^ responses in both pericytes and vascular smooth muscle cells (Fig. 3). This blunting of vasomotor activity was paralleled by suppressed mural cell Ca^2+^ signals, indicating inhibition of myogenic tone. In contrast, endothelial cell Ca^2+^ imaging revealed the opposite effect: high-viscosity perfusate enhanced flow-induced Ca^2+^ transients, implicating shear-stress-sensitive endothelial pathways. The response resembled that observed in studies of the Piezo1 channel, a mechanosensitive channel activated by shear stress and linked to eNOS activation (14). Consistent with this, NOS inhibition with L-NAME restored mural cell Ca^2+^ influx under high-viscosity conditions, confirming that eNOS-derived NO suppresses pressure-induced tone. Together, these results support a model in which elevated plasma viscosity increases endothelial shear stress, activates Piezo1 and related mechanosensors, and drives NO release that diffuses to adjacent mural cells, suppressing their Ca^2+^ activity and vasoconstriction. This establishes an autoregulatory feedback loop between endothelial cells and pericytes/vSMCs in regulating capillary perfusion.

**Figure 4.**
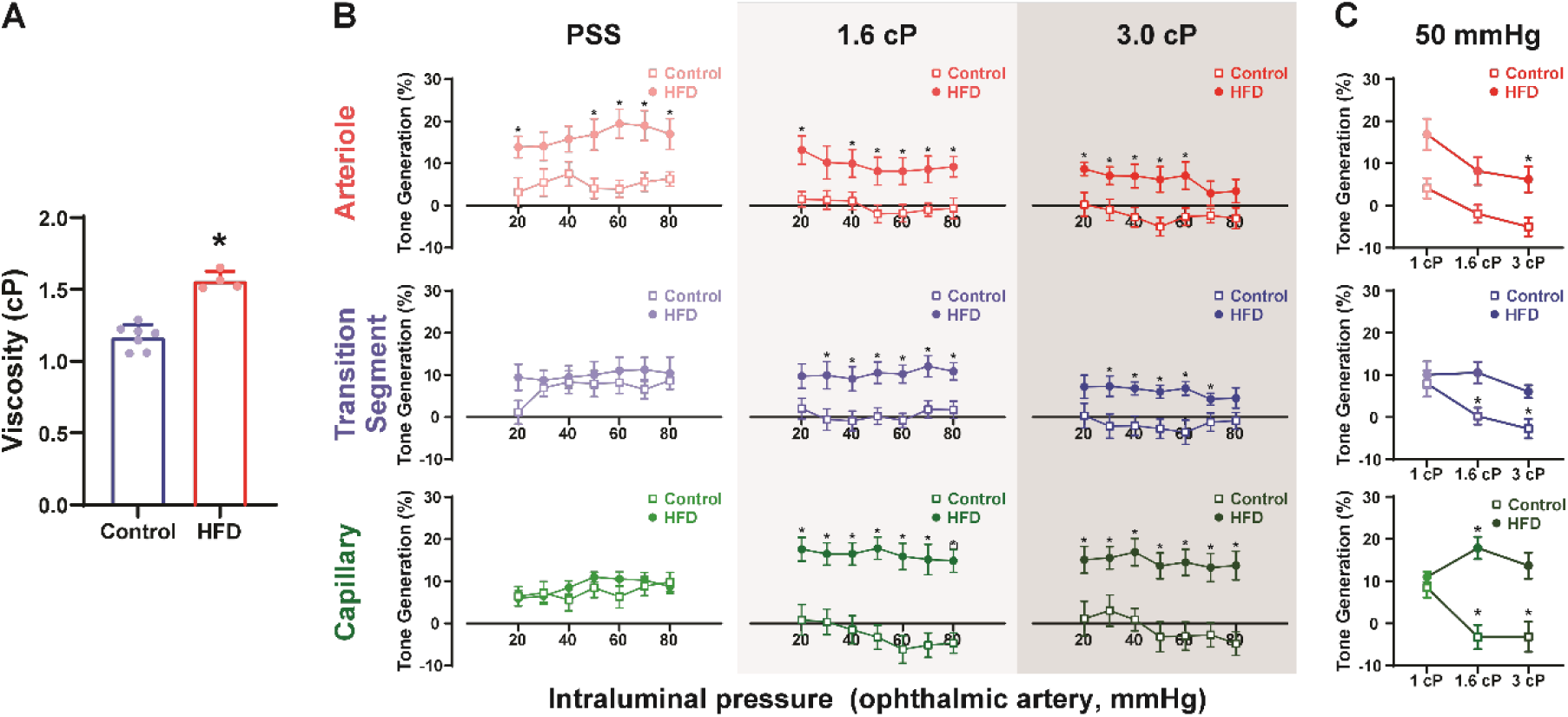
High Viscosity Inhibits Pressure-induced Tone: **A:** Summary of plasma viscosity measurements from 3-month-old mice on either normal chow (WT) or a high-fat diet (HFD). Data presented with means ± SEM (*p ≤ 0.05 vs control); and N = 4-7 mice. **B:** Summary data of the pressure-dependent diameter changes in arterioles (top, red), transition segments (middle, blue), and capillaries (bottom, green) under three perfusate conditions: PSS, 1.6 cP, and 3.0 cP. Diameter responses to step increases in intraluminal pressure (20–80 mmHg) for WT (open symbols) and HFD (filled symbols). Data presented with means ± SEM (*p ≤ 0.05 vs control); two-way repeated-measure ANOVA with Tukey’s multiple comparison test; n=7-20; and N = 3-5 mice. **C:** Summary of tone generation at 50 mmHg, showing that mice fed a high-fat diet exhibit impaired viscosity-dependent loss of tone in transition-segment and capillary vessels. Data presented with means ± SEM; *p ≤ 0.05 vs 1 cP; two-way repeated-measure ANOVA with Tukey’s multiple comparison test; n=10-28; and N = 3-5 mice.

**Figure 5.**
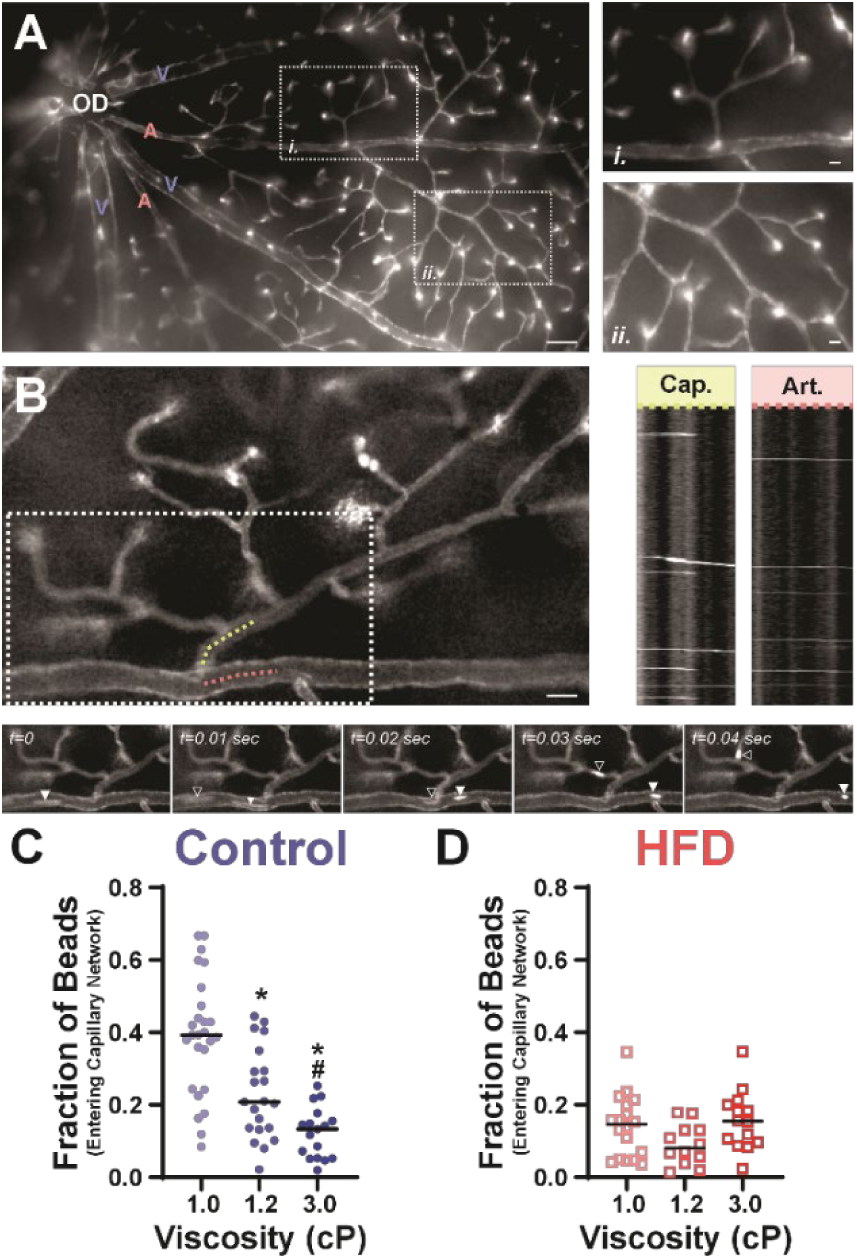
Increased Intraluminal Viscosity Reduces Capillary Entry of Flowing Microspheres. **A:** Representative image of fluorescent isolectin staining showing the large-scale architecture of the retinal vasculature. Insets: **(*i*)** representative image of a lateral/orthogonal branch pattern and **(*ii*)** representative image of a dichotomous branching pattern. **B:** Representative image (top left), representative time-series frames (bottom), and representative kymographs (top right) illustrating microsphere tracking across vascular segments. Dotted lines indicate regions used for kymograph generation in an arteriole (red) and a transition segment (yellow). Time-series frames show one microsphere entering the capillary network (open arrowhead) and another remaining within the arteriole (closed arrowhead). Kymographs highlight microspheres diverging into capillaries (yellow) or continuing along the arteriole (red). **C:** Quantification of the fraction of microspheres entering capillary segments versus remaining in the arteriole in control mice. Controls exhibit high variability and greater capillary entry at 1 cP, which declines at 1.2 cP and 3.0 cP. **D:** Mice fed a high-fat diet (HFD) show reduced capillary entry across all viscosities, indicating impaired microsphere distribution into the capillary network. Data are presented as mean ± SEM; *p ≤ 0.05 vs PSS and #p ≤ 0.05 vs 1.2 cP; one-way ANOVA with Tukey’s multiple-comparison test; n = 14-20 capillary networks; and N = 6 mice.

### High Fat Diet-induced Vascular Remodeling

The present findings also have implications for how chronic dietary changes, such as high-fat intake, may alter microvascular structure and function. High-fat diets have been shown to impair endothelial function, increase vascular stiffness, and reduce myogenic tone in arterioles. As a result, arterioles remodel hypertrophically by increasing elastin and collagen to increase stiffness, as well as phenotypic switching to macrophage/fibroblast-like cells following cholesterol exposure (27, 28). To examine the effects on the capillary microcirculation, we fed mice a high-fat diet and found that the resulting increase in plasma viscosity enhanced baseline arteriolar tone but had no effect on capillary pericytes. Unlike arterioles, we did not observe an increase in pressure-induced tone generation in either ensheathing or thin-stranded pericytes, suggesting an absence of remodeling in these cells. Others have observed increases in NG2 and α-SMA-positive ensheathing pericytes within the transition segment after reduction of Notch3 in Col4a1^+/G498V^ mice, possibly due to an increase in pressure from a lack of SMCs in the arteriole (29). Rather than evidence of mural cell remodeling, we observed an altered capillary endothelial response, in which viscosity-dependent increases in shear stress failed to reduce pressure-induced tone in capillary pericytes, indicating a blunted mechanosensory response. These changes suggest that elevated plasma viscosity, driven by dietary factors, may reduce the predictability and efficiency of blood flow entering the capillary network.

This effect may be exacerbated *in vivo* by the presence of RBCs, which, under slow-flow conditions, increase whole-blood viscosity and worsen perfusion deficits. In the context of high-fat diets or metabolic syndrome, increased plasma proteins, lipids, and altered hematocrit could further raise blood viscosity, leading to chronic hypoperfusion and maladaptive vascular remodeling. These findings underscore how diet-induced changes in blood rheology may contribute to microvascular dysfunction and potentially vascular rarefaction over time.

### Clinical Impact of Plasma Viscosity on Cardiovascular Disease

The clinical relevance of these findings is underscored by conditions such as hyperviscosity syndrome (30), where excessive blood viscosity causes systemic hypoperfusion, with neurological symptoms including headache, dizziness, seizures, and stroke (30). Elevated viscosity is also associated with cardio-cerebrovascular diseases such as atherosclerosis and ischemic stroke, which may accelerate Alzheimer’s disease by worsening vasculopathy (16, 17, 31–33). Reduced perfusion of metabolically active neurons may contribute to cognitive decline through chronic hypoxia. Importantly, blood viscosity is strongly influenced by metabolic health (21, 34–36). Nutrient composition, plasma proteins, and circulating lipids all affect viscosity, helping explain why individuals with metabolic syndrome, which affects ∼1 in 3 adults in the U.S., exhibit elevated viscosity, higher blood pressure, and greater risk for diabetes, heart disease, and stroke (9, 37). Despite these associations, blood viscosity remains an underappreciated factor in cardiovascular risk assessment.

## Conclusion

This study directly demonstrates that increased plasma viscosity alters microcirculatory dynamics by delaying capillary filling, suppressing pressure-induced tone, and enhancing endothelial shear stress. These changes reflect a shift in flow regulation from mural cell constriction to endothelial NO-mediated feedback. While RBCs were absent here, their presence in vivo would likely amplify viscosity-induced impairments due to shear-thinning dynamics and capillary deformation. Collectively, these findings highlight viscosity as a key but underrecognized determinant of vascular health and underscore the need for further work to dissect its role in microvascular disease and vascular injury.

## Acknowledgments

This study was supported by grants from the BrightFocus Foundation (M2022010N to A.L.G.), NIH (NIA R01AG081935, NHLBI R01NHLBI, NHLBI K01HL138215, and NIGMS P20GM130459 to A.L.G.). The Transgenic Genotyping and Phenotyping Core and the High Spatial and Temporal Resolution Imaging Core at the Centers of Biomedical Research Excellence (COBRE) Center for Molecular and Cellular Signaling in the Cardiovascular System, University of Nevada, Reno are maintained by grants from NIH/NIGMS (P20GM130459 Sub#5451 and P20GM130459 Sub#5452). We thank E. Franco, P. Voss, M. Dagda, and B. Harper for technical assistance and animal care, A. Aupetit for editorial assistance, and R. Allen for microscopy assistance.

## Author contributions

C.E.F. and A.L.G. designed research; C.E.F. performed research; C.E.F. and A.L.G. analyzed data; and C.E.F. and A.L.G. wrote the paper.

## References

1. S. Grubb et al., Precapillary sphincters maintain perfusion in the cerebral cortex. Nature Communications 11, 395 (2020).

2. P. D. Harris, D. E. Longnecker, Significance of precapillary sphincter activity for microcirculatory function. Microvasc Res 3, 385–395 (1971).

3. P. Thakore et al., Brain endothelial cell trpa1 channels initiate neurovascular coupling. eLife 10, 1–84 (2021).

4. T. A. Longden et al., Capillary K+-sensing initiates retrograde hyperpolarization to locally increase cerebral blood flow. Nature neuroscience 20, 717–717 (2017).

5. J. L. Taylor et al., Functionally linked potassium channel activity in cerebral endothelial and smooth muscle cells is compromised in Alzheimer’s disease. Proceedings of the National Academy of Sciences 119 (2022).

6. N. R. Klug et al., Intraluminal pressure elevates intracellular calcium and contracts CNS pericytes: Role of voltage-dependent calcium channels. Proceedings of the National Academy of Sciences 120, e2216421120–e2216421120 (2023).

7. H. R. Ferris et al., Increased luminal pressure in brain capillaries drives TRPC3-dependent depolarization and constriction of transitional pericytes. Science Signaling 18, eads1903 (2025).

8. A. L. Gonzales et al., Contractile pericytes determine the direction of blood flow at capillary junctions. Proc Natl Acad Sci U S A 117, 27022–27033 (2020).

9. G. Sloop, R. E. Holsworth, J. J. Weidman, J. A. St Cyr, The role of chronic hyperviscosity in vascular disease. Therapeutic Advances in Cardiovascular Disease 9, 19–25 (2014).

10. E. Nader et al., Blood Rheology: Key Parameters, Impact on Blood Flow, Role in Sickle Cell Disease and Effects of Exercise. Front Physiol 10, 1329 (2019).

11. A. M. Eltanahy, A. Aupetit, E. D. Buhr, R. N. Van Gelder, A. L. Gonzales, Light-sensitive Ca^2+^ signaling in the mammalian choroid. Proceedings of the National Academy of Sciences 121, e2418429121 (2024).

12. O. F. Harraz, N. R. Klug, A. J. Senatore, D. C. Hill-Eubanks, M. T. Nelson, Piezo1 Is a Mechanosensor Channel in Central Nervous System Capillaries. Circ Res 130, 1531–1546 (2022).

13. A. Mughal, M. T. Nelson, D. Hill-Eubanks, The post-arteriole transitional zone: a specialized capillary region that regulates blood flow within the CNS microvasculature. The Journal of Physiology 601, 889–901 (2023).

14. O. F. Harraz, N. R. Klug, A. J. Senatore, D. C. Hill-Eubanks, M. T. Nelson, Piezo1 Is a Mechanosensor Channel in Central Nervous System Capillaries. Circulation Research 130, 1531–1546 (2022).

15. S. Wang et al., Endothelial cation channel PIEZO1 controls blood pressure by mediating flow-induced ATP release. Journal of Clinical Investigation 126, 4527–4536 (2016).

16. G. Chen et al., Regulation of blood viscosity in disease prevention and treatment. Chinese Science Bulletin 57, 1946–1952 (2012).

17. S. Aras et al., Plasma Viscosity: Is a Biomarker for the Differential Diagnosis of Alzheimer’s Disease and Vascular Dementia? American Journal of Alzheimer’s Disease & Other Dementias® 28, 62–68 (2012).

18. A. L. Gonzales et al., A PLCγ1-dependent, force-sensitive signaling network in the myogenic constriction of cerebral arteries. Science signaling 7, ra49–ra49 (2014).

19. S. Earley, B. J. Waldron, J. E. Brayden, Critical Role for Transient Receptor Potential Channel TRPM4 in Myogenic Constriction of Cerebral Arteries. Circulation Research 95, 922–929 (2004).

20. M. M. Y Schnitzler et al., Gq-coupled receptors as mechanosensors mediating myogenic vasoconstriction. EMBO Journal 27, 3092–3103 (2008).

21. I. Cicha, Y. Suzuki, N. Tateishi, N. Maeda, Effects of dietary triglycerides on rheological properties of human red blood cells (abstract). Clinical Hemorheology and Microcirculation 30, 301–305 (2004).

22. K. L. Sweazea, B. R. Walker, Impaired myogenic tone in mesenteric arteries from overweight rats. Nutrition & Metabolism 9, 18 (2012).

23. Victor V. Lima et al., High-fat diet increases O-GlcNAc levels in cerebral arteries: a link to vascular dysfunction associated with hyperlipidaemia/obesity? Clinical Science 130, 871–880 (2016).

24. S. M. Sargent, et al. (2023) Endothelial structure contributes to heterogeneity in brain capillary diameter. in bioRxiv: the preprint server for biology.

25. A. Borovoi, E. Naats, U. Oppel, Scattering of light by a red blood cell. Journal of Biomedical Optics 3 (1998).

26. B. Y. S. Vázquez, Blood pressure and blood viscosity are not correlated in normal healthy subjects. Vascular health and risk management 8, 1–6 (2012).

27. I. A. M. Brown et al., Vascular Smooth Muscle Remodeling in Conductive and Resistance Arteries in Hypertension. Arterioscler Thromb Vasc Biol 38, 1969–1985 (2018).

28. A. Chattopadhyay et al., Cholesterol-Induced Phenotypic Modulation of Smooth Muscle Cells to Macrophage/Fibroblast–like Cells Is Driven by an Unfolded Protein Response. *Arteriosclerosis*, Thrombosis, and Vascular Biology 41, 302–316 (2021).

29. J. Ratelade et al., Reducing Hypermuscularization of the Transitional Segment Between Arterioles and Capillaries Protects Against Spontaneous Intracerebral Hemorrhage. Circulation 141, 2078–2094 (2020).

30. A. Perez Rogers, M. Estes, “Hyperviscosity Syndrome”. (2023).

31. S. H. Chew, M. Meighan Tomic, A. T. W. Cheung, Alzheimer’s disease: More than amyloid. Clinical Hemorheology and Microcirculation 46, 69–73 (2010).

32. E. Cecchi et al., Hyperviscosity as a Possible Risk Factor for Cerebral Ischemic Complications in Atrial Fibrillation Patients. The American Journal of Cardiology 97, 1745–1748 (2006).

33. A. J. Lee et al., Blood viscosity and elevated carotid intima-media thickness in men and women: the Edinburgh Artery Study. Circulation 97, 1467–1473 (1998).

34. D. Destiana, I. S. Timan, The relationship between hypercholesterolemia as a risk factor for stroke and blood viscosity measured using Digital Microcapillary ®. 10.1088/1742-6596/1073/4/042045, 42045–42045 (2018).

35. H. Naghedi-Baghdar et al., Effect of diet on blood viscosity in healthy humans: a systematic review. Electronic Physician 10, 6563–6563 (2018).

36. D.-d. Pan et al., [Effect of improper diets on blood viscosity in SD rats in high-salt and fat diet and alcohol abuse simulation model]. Zhongguo Zhong yao za zhi = Zhongguo zhongyao zazhi = China journal of Chinese materia medica 40, 1560–1564 (2015).

37. C. Irace, F. Scavelli, C. Carallo, R. Serra, A. Gnasso, Plasma and blood viscosity in metabolic syndrome. *Nutrition*, Metabolism and Cardiovascular Diseases 19, 476–480 (2009).

